# A scientometric overview of CORD-19

**DOI:** 10.1101/2020.04.20.046144

**Authors:** Giovanni Colavizza, Rodrigo Costas, Vincent A. Traag, Nees Jan van Eck, Thed van Leeuwen, Ludo Waltman

## Abstract

As the COVID-19 pandemic unfolds, researchers from all disciplines are coming together and contributing their expertise. CORD-19, a dataset of COVID-19 and coronavirus publications, has been made available alongside calls to help mine the information it contains and to create tools to search it more effectively. We analyse the delineation of the publications included in CORD-19 from a scientometric perspective. Based on a comparison to the Web of Science database, we find that CORD-19 provides an almost complete coverage of research on COVID-19 and coronaviruses. CORD-19 contains not only research that deals directly with COVID-19 and coronaviruses, but also research on viruses in general. Publications from CORD-19 focus mostly on a few well-defined research areas, in particular: coronaviruses (primarily SARS-CoV, MERS-CoV and SARS-CoV-2); public health and viral epidemics; molecular biology of viruses; influenza and other families of viruses; immunology and antivirals; clinical medicine. CORD-19 publications that appeared in 2020, especially editorials and letters, are disproportionately popular on social media. While we fully endorse the CORD-19 initiative, it is important to be aware that CORD-19 extends beyond research on COVID-19 and coronaviruses.

## Introduction

The COVID-19 pandemic is attracting the attention of the global scientific community. Biomedical research on the virus and on the management of the crisis from an epidemiological and healthcare point of view has full priority. Furthermore, many research communities, funding agencies and third-parties are taking action to support the fight against the pandemic with their own expertise and resources. Multidisciplinary open collaboration will prove instrumental to fight the current and future pandemics [6, 5].

Several initiatives have been taken to share COVID-19-related scientific research as openly as possible, from public, private and non-profit organisations. It has been readily recognized that health crises are also information crises [39, 11, 31, 20]. For example, the World Health Organization maintains a list of relevant research updated daily [41], as well as a portal to provide information to the public [2], similar to the the European Commission [3]. Publishers are opening access to relevant publications, and some open-access publishers have lifted publishing fees for the same reason.

Another initiative is the release of the COVID-19 Open Research Dataset (CORD-19) [38]. CORD-19 is a growing, daily-updated dataset of COVID-19 publications, capturing new as well as past research on “COVID-19 and the coronavirus family of viruses for use by the global research community.”^1^ The reason to release this dataset is “to mobilize researchers to apply recent advances in natural language processing to generate new insights in support of the fight against this infectious disease.” The initiative has the backing of the US White House [1] and is a partnership of several institutions including the Allen Institute for Artificial Intelligence, the Chan Zuckerberg Initiative, Georgetown University’s Center for Security and Emerging Technology, Microsoft Research, the National Library of Medicine of the National Institutes of Health, and Unpaywall. CORD-19 is released together with a set of challenges hosted by Kaggle, mainly focused on automatically extracting structured and actionable information from such a large set of publications. The release of this dataset is a positive call for action directed towards the natural language processing, machine learning and related research communities.^2^

In order to contribute to an informed use of the CORD-19 dataset, we present a scientometric analysis of its contents. We first consider the subject delineation of CORD-19. Field or subject delineation is a complex task, which usually benefits from an interplay of information retrieval, or search-based approaches, and bibliometric mapping [19, 42, 24]. Based on a comparison to the Web of Science database, we find that CORD-19 provides an almost complete coverage of research on COVID-19 and coronaviruses. We also show that CORD-19 covers other research topics beyond COVID-19 and coronaviruses.

We then provide a detailed analysis of the research covered by CORD-19, enriching the dataset with data from Dimensions [18] and using both citation analysis and text analysis. We show that CORD-19 broadly focuses on biomedical research on viruses and related health issues. The articles it covers are quite heterogeneous. CORD-19 contains a core of research directly on COVID-19 and coronaviruses, but in addition it contains many articles on related yet distinct streams of virus research, such as on influenza, molecular biology and public health. Secondly, we identify three broad periods in the accumulation of literature in CORD-19: a pre-SARS (2003) period, a post-SARS period and the current pandemic (2020). We also present a brief analysis of Altmetric data [30, 27] related to papers in CORD-19. We find that papers published in 2020 are disproportionately represented in social media and news by Altmetric indicators, especially on Twitter. We also observe that non-research articles, such as editorials and letters, are particularly popular on social media. We conclude by emphasizing the importance of being aware of the broad coverage of CORD-19, which extends beyond research on COVID-19 and coronaviruses. We also propose some directions for future work.

To facilitate the analysis and use of the CORD-19 dataset, we release our code (see Appendix). In combination with access to Dimensions, Altmetric, Web of Science and Twitter, our results can be replicated. Finally, we underline that we have no specific biomedical expertise. We invite experts to improve the interpretation of our results.

## The CORD-19 dataset

The CORD-19 dataset [38] is being updated on a daily basis. We use the version of the dataset from July 1, 2020. This version of CORD-19 contains 169, 821 articles, of which 77, 777 are equipped with full text. The dataset collects publications from the following sources:

- Medline and PubMed’s PMC open access corpus, via the query: `“COVID-19” OR Coronavirus OR “Corona virus”` `OR “2019-nCoV” OR “SARS-CoV” OR “MERS-CoV”` `OR “Severe Acute Respiratory Syndrome” OR` `“Middle East Respiratory Syndrome”.`
- COVID-19 research articles from the WHO database [4].
- arXiv, bioRxiv and medRxiv pre-prints using the same query as in PMC.

A relatively small number of publications prior to the SARS 2003 outbreak is followed by a steady increase in the number of publications up to 2020, when the growth accelerated substantially (Figure 2). The top 20 sources (i.e., journals and pre-print servers) by number of publications are given in Figure 1, highlighting how CORD-19 is composed of publications from varied sources. Pre-print servers, the Journal of Virology and PLOS ONE stand out. The main contributors to the dataset are Medline, PMC and Elsevier (Figure A.1b), which have made a large part of their relevant literature available. The availability of full texts is relatively high, but not yet complete, and is proportionally stable over time (Figure A.1a). This is because, even though most publishers and journals have opened up their publications, some have not yet done so. In particular, the Journal of Virology, Nature, Science, The Lancet and BMJ provided no or only few articles in full text at the time of our analysis (July 2020).

**Figure 1:**
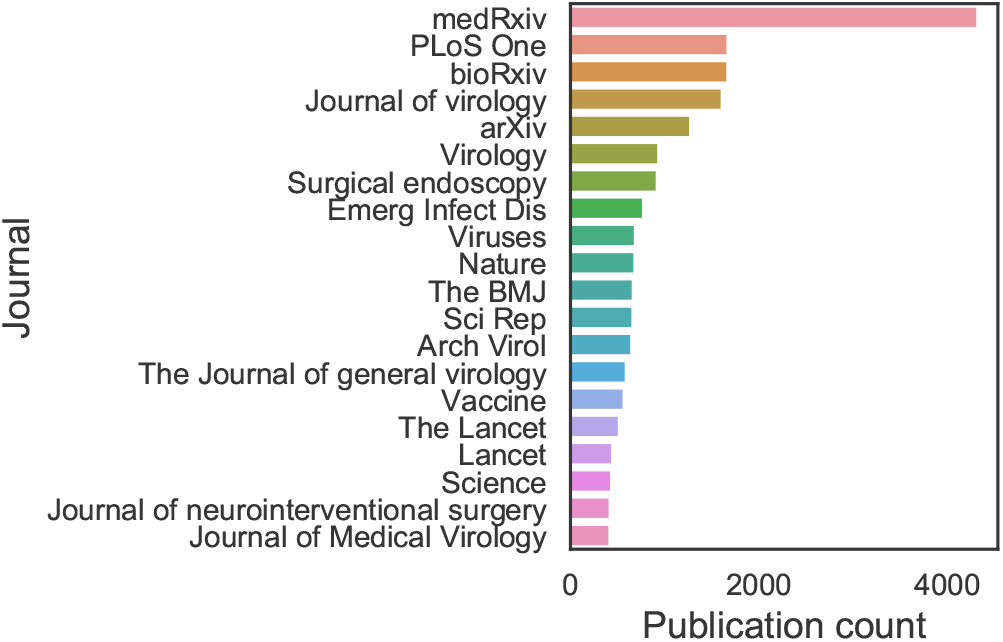
Top 20 journals and pre-print servers by number of CORD-19 articles.

**Figure 2:**
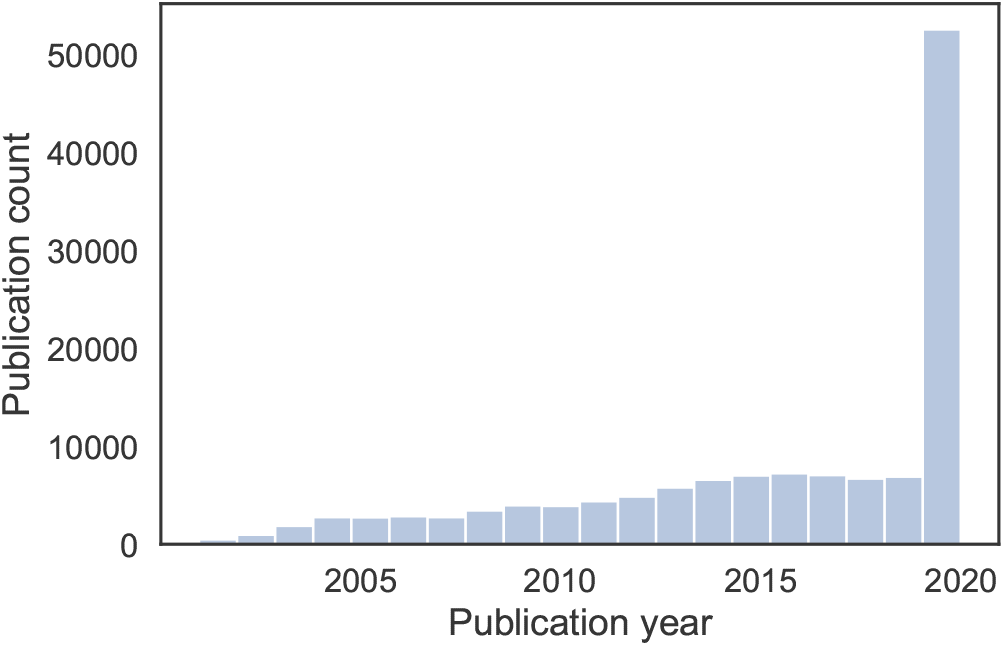
Publication years of CORD-19 papers.

Using Dimensions data, we study the number of citations to papers in CORD-19 (Figure A.2). The literature in CORD-19 gained more attention from the early 2000s, and some publications from 2020 are accumulating citations at an extreme rate. The most cited articles from 2020 include: “Clinical features of patients infected with 2019 novel coronavirus in Wuhan, China” by Huang et al. (2020, The Lancet, also the most cited overall), “Clinical Characteristics of Coronavirus Disease 2019 in China” by Guan et al. (2020, The New England Journal of Medicine), “A Novel Coronavirus from Patients with Pneumonia in China, 2019” by Zhu et al. (2020, The New England Journal of Medicine) and “Clinical Characteristics of 138 Hospitalized Patients With 2019 Novel Coronavirus–Infected Pneumonia in Wuhan, China” by Wang et al. (2020, JAMA). The Lancet is the most cited journal in the dataset in terms of the absolute number of citations received from papers in CORD-19, based on Dimensions data. It is followed by PLOS ONE and Nature. Lastly, we consider the Fields of Research (FOR) categories provided by Dimensions, which cover all areas of research from the Australian and New Zealand Standard Research Classification (ANZSRC).^3^ While about 30, 000 articles have no classification, we find that, as expected, the Medical and Health Sciences cover most ground, followed by Biological Sciences (Figure A.3). The second level FOR classification shows the presence of several sub-areas of these two top level fields, Clinical Sciences being the largest. More details and plots are provided in the accompanying repository.

## Delineation of CORD-19

We now analyse how the literature on COVID-19 and coronaviruses is delineated in CORD-19. We first examine whether the delineation might be too broad, meaning that CORD-19 would include publications that are of limited relevance for coronavirus research. We then examine whether the delineation might be too narrow, meaning that relevant publications would not be included in CORD-19. We focus on publications from the time period 1980–2019. We do not consider publications from 2020, because most of these publications are not yet available in the Web of Science (WoS) database used in our analysis.

Figure 3 shows the annual number of publications in CORD-19 as well as the annual number of publications in a subset of CORD-19 that we refer to as ‘CORD-19-strict’. CORD-19-strict consists of all publications in CORD-19 that we have identified by applying the CORD-19 search query to the titles and abstracts of publications but not to their full texts. Hence, publications for which the CORD-19 search query yields a match only in the full text and not in the title and abstract are not included in CORD-19-strict. These publications probably do not directly focus on COVID-19 and coronaviruses, even though they may still be related to these topics in some way. If these publications had been directly focused on COVID-19 and coronaviruses, it can be expected that applying the CORD-19 search query to the title and abstract would have resulted in a match.

**Figure 3:**
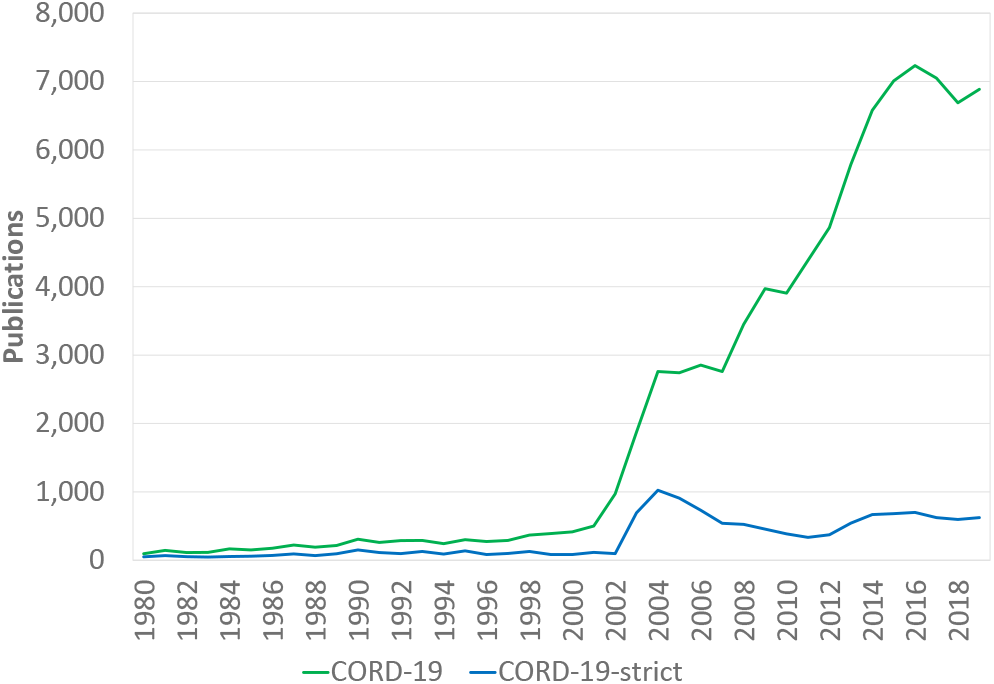
Annual number of publications in CORD-19 and CORD-19-strict.

As can be seen in Figure 3, the number of CORD-19 publications started to increase around 2002 and 2003, which coincides with the 2002–2004 SARS outbreak. Looking at CORD-19-strict, a second increase took place after 2012, following the 2012 MERS outbreak. More significantly for our analysis, Figure 3 shows that CORD-19-strict includes only a relatively small subset of all publications in CORD-19. This suggests that CORD-19 includes many publications that are not directly focused on COVID-19 and coronaviruses research. The inclusion of these publications in CORD-19 does no harm and may be useful for a broader perspective on coronoviruses research. However, it is important for users of CORD-19 to be aware that the dataset includes a large number of publications whose relevance for COVID-19 and coronaviruses research needs a more careful assessment, and some of which may be of limited relevance.

We now take an opposite perspective, aiming to find out whether relevant publications may be missing in CORD-19. To do so, we use the CWTS in-house version of the WoS database [7].^4^ We consider the following WoS citation indices: Science Citation Index Expanded, Social Sciences Citation Index, Arts & Humanities Citation Index, and Conference Proceedings Citation Index. We use the CORD-19 search query to identify relevant publications in WoS. The search query is applied to the titles, abstracts, and keywords of publications, where we note that abstracts and keywords are available only for publications from 1991 onward. We refer to the resulting set of publications as ‘WoS-strict’, where ‘strict’ indicates that publications have been identified in a similar way as in CORD-19-strict, so without searching in the full texts of publications.

Figure 4 shows the annual number of publications in WoS-strict. In addition, the figure shows the annual number of publications in WoS-strict that are also included in CORD-19, using DOIs and PubMed IDs to match publications in the two datasets. We see that almost all publications in WoS-strict can also be found in CORD-19. This indicates that CORD-19 provides an almost complete coverage of publications that have a clear relevance for research on COVID-19 and coronaviruses, at least when compared to WoS. Importantly, in a comparison of CORD-19 and WoS that we performed a few months previous to the one we report here (discussed in an earlier version of the present paper [12]), we found substantial gaps in the coverage of the coronavirus literature by CORD-19 In a span of a few months, it appears that most gaps in the coverage of CORD-19 have been filled. This testifies to the efforts made by the CORD-19 team to expand the coverage of the dataset [21].

**Figure 4:**
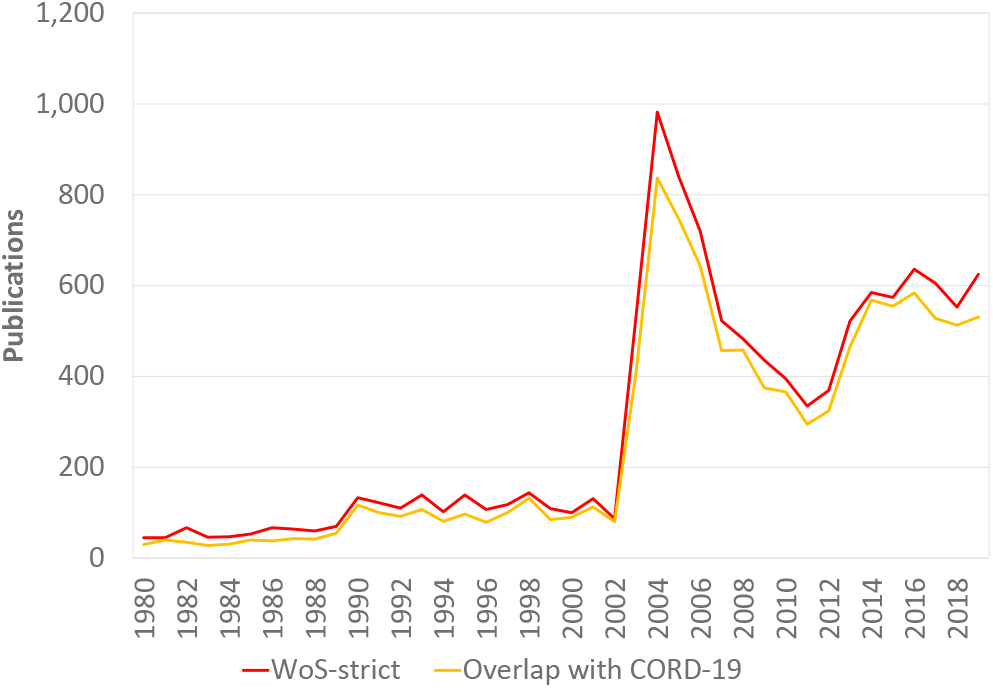
Annual number of publications in WoS-strict and the overlap with CORD-19.

## Analysis of CORD-19

We conducted the following analyses on the CORD-19 dataset: a term map and a topic modelling analysis based on titles and abstracts of publications in CORD-19; a citation network analysis using Dimensions citation data; an altmetrics analysis using Altmetric data. The purpose of the former two analyses is to cluster CORD-19 publications and further clarify the contents of the dataset. For our analyses, we focus on 140, 302 articles out of the 169, 821 available in CORD-19, filtering out duplicates and articles without at least one identifier among DOI, PubMed ID, PubMed Central ID or arXiv ID.

### Term map

To get an accessible high-level overview of the contents of CORD-19, we used VOSviewer [33] to create a so-called term map of the publications in this dataset. Titles and abstracts of publications were concatenated into a single string and were then provided as input to VOSviewer. Using the text mining algorithms of VOSviewer, we identified the 2, 115 most relevant terms in the titles and abstracts. A term is defined as a sequence of nouns and adjectives ending with a noun. Only terms occurring in at least 130 publications were considered. Plural terms were converted to singular. For each pair of terms, VOSviewer counted the number of publications in which both terms occur in the title or abstract. In this way, a co-occurrence network was obtained, indicating for each pair of terms the number of publications in which they occur together.

Figure 5 presents a visualisation of this co-occurrence network. The visualisation, referred to as a term map, shows the 2, 115 terms included in the network. The size of a term reflects the number of publications in which the term occurs. The proximity of two terms in the map approximately indicates the relatedness of the terms based on their number of co-occurrences. In general, the closer two terms are located to each other, the stronger they are related. This means that groups of terms located closely together in the map usually can be interpreted as topics. The horizontal and vertical axes have no special meaning. Labels are shown only for a selection of the terms in the term map. In order to avoid overlapping labels, for many of the less prominent terms no label is shown. The term map can also be explored interactively online (https://bit.do/term_map_cord19_20200701). The labels of the less prominent terms can then be made visible by zooming in on specific areas in the map.

**Figure 5:**
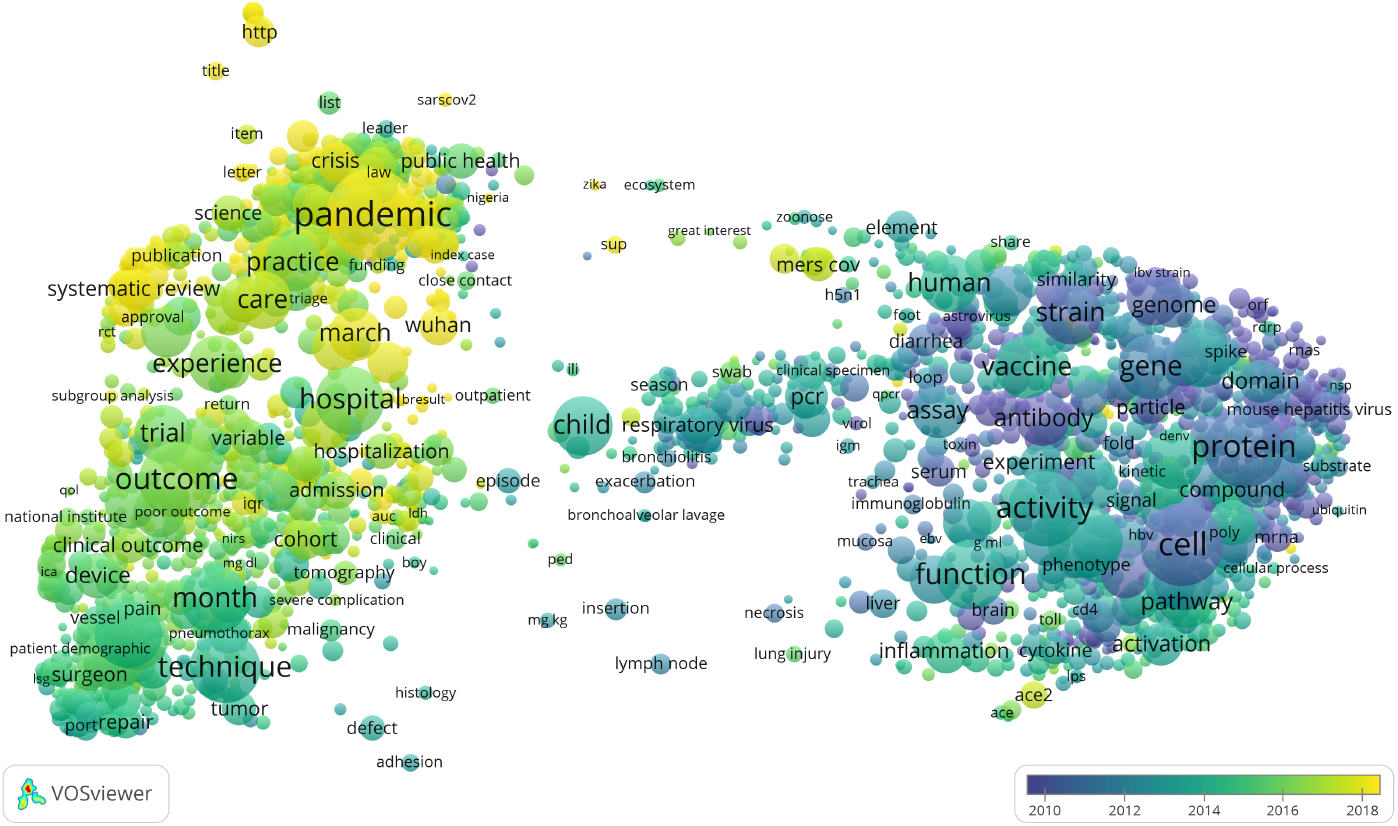
Term map highlighting temporal trends in the contents of CORD-19. Also compare with the topic words in the Appendix.

Colors can be used to present additional information in the term map. In Figure 5, for instance, the color of a term reflects the average publication year of the publications in which the term occurs. In this way, temporal trends are made visible. The term map shows a clear divide between biologically focused topics in the right area in the map and clinically focused topics and health research in the left area (for similar findings, see [34]). As shown by the colors of the terms, the topics in the left area in the map received a lot of attention in more recent years, while the topics in the right area received attention mostly in earlier years. Below, we use our term map to illustrate some of the analyses we present.

### Topic modelling

We conducted a topic modelling analysis, making use of the titles and abstracts of CORD-19 publications by concatenating them into a single string, similarly to what we did for the term map. Of the 140, 302 articles, 31, 055 have no abstract. Topic modelling is a technique to learn ‘topics’ from a corpus of documents. A topic is defined as a probability distribution over a vocabulary, where words with a high probability for the same topic tend to co-occur frequently in the same documents [8, 40, 23]. We applied a pre-processing pipeline using ScispaCy’s `en_core_sci_md_model` [26] to convert each publication’s title and abstract into a bag-of-words representation. This included the following steps: entity detection and inclusion in the bag-of-words for entities consisting of at least two tokens; lemmatisation; removal of (isolated) punctuation and stopwords; inclusion of frequent bigrams.

We started by training a Latent Dirichlet Allocation (LDA) model [10], using gensim’s implementation [28] and 15 topics. From a topic coherence analysis [25], we found 15 to 25 to be a good value for the number of topics. We decided to remain on the lower end to facilitate the interpretation of the results. We further filtered out words that appear in fewer than ten or in more than half of the documents. Lastly, we verified our results using a Correlated Topic Model [9],^5^ which reduces to a Structural Topic Model without covariates [29]. This model explicitly captures topic correlations. It confirmed the topic structure we found with LDA, as well as the topics’ temporal unfolding. More details are provided in the accompanying repository. We report results of the LDA model in what follows.

The top 20 words per topic are given in the Appendix. In Figure A.4, we show the yearly topic intensity, from 1980 to 2020. Three periods in the relative focus of the literature seem to emerge. Pre-SARS research mainly focused on molecular biology and immunology (topics 3, 8, 11 and 12). Subsequently, due primarily to the SARS and MERS outbreaks (topic 5), the literature broadened its focus to include topics such as epidemics (topics 9 and 10), public health (topics 0, 4 and 13) and clinical medicine (topics 2, 6, 7 and 14). Lastly, in 2020 the literature is also heavily focused on the COVID-19 outbreak, its ongoing management and consequences (topic 1). From this first analysis, we observe how coronavirus research seems to be produced in bursts following outbreaks instead of following a more steady progress.

In order to analyse the contents of CORD-19 at a higher level of granularity, we grouped the identified topics into *general topics*. We used agglomerative clustering based on the Jensen-Shannon distances among topic-word distributions to inform the grouping of topics (see the accompanying repository for more details). To be sure, the resulting grouping is a simplification of the actual publication contents. It is also worth considering that topics overlap substantially. The following general topics were identified:

- “Clinical medicine”: topics 2, 6, 7, 14;
- “Coronavirus outbreaks” (SARS, MERS, COVID-19): topics 1, 5;
- “Epidemics”: topics 9, 10;
- “Immunology”: topics 8, 12;
- “Molecular biology”: topics 3, 11;
- “Public health”: topics 0, 4, 13.

CORD-19 is dominated by literature on coronavirus outbreaks, and to a lesser degree public health, epidemics and clinical medicine. There are fewer publications focusing on molecular biology and immunology. Figure 6 shows the relative (average per paper) and absolute (total over all papers) topic intensities of the general topics. The temporal development of the contents of CORD-19 discussed above is largely confirmed. The 2003 SARS outbreak generated a shift associated with a rise in publications on coronavirus epidemics, which is happening again, at a much larger scale, in 2020 for COVID-19. Molecular biology and immunology research, on the other hand, have lost ground in relative terms.

**Figure 6:**
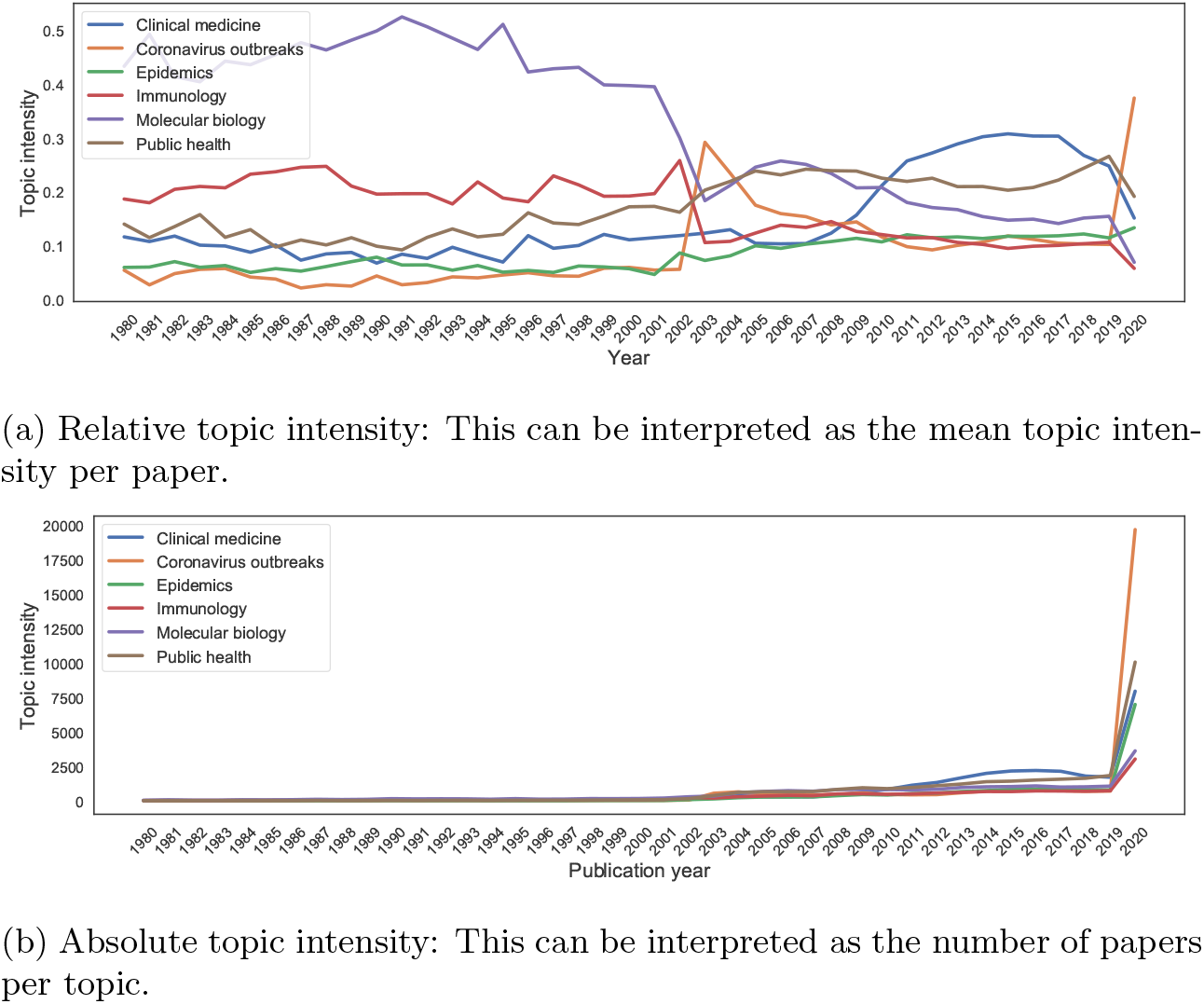
Intensity of the general topics over time.

### Citation clusters

The previous two analyses were based on textual data. We now turn to a view based on citations to characterise the CORD-19 dataset.

We constructed a citation network based on the references included in all papers in CORD-19, as provided by Dimensions. We considered not only references to papers in CORD-19, but also all other references in CORD-19 papers. These ‘external’ references provide additional information regarding the structure of CORD-19. For example, two papers may not be immediately connected via papers in the CORD-19 dataset, but they may have common references external to CORD-19. We considered only the giant weakly connected component, which amounted to 1, 844, 831 publications and 3, 974, 851 citation links, of which 104, 508 publications belong to the CORD-19 dataset. This is the citation network that we work with in the remainder of this section.

We clustered the citation network using the Leiden algorithm^6^ [32]. Each citation link was weighted as 1*/k_i_*, where *k_i_* is the number of references of publication *i*. The inclusion of the ‘external’ references in the context of clustering is also known as the extended direct citation approach [37]. In this approach, the publications contained in CORD-19 are weighted with a so-called node weight of 1, while the ‘external’ publications are weighted with a weight of 0 (see [37] for more details). We clustered the citation network in a hierarchical fashion. At the most detailed level, the citation network was clustered using a resolution of 2 10^*−*5^. We aggregated the citation network based on this clustering, and then clustered the resulting aggregated citation network at a resolution of 1·10^*−*5^. We refer to the former clustering as the bottom level clustering, and to the latter as the top level clustering. The ‘external’ publications help cluster publications from CORD-19 more accurately. In what follows, we focus only on the 104, 508 publications included in CORD-19.

The clustering at the bottom level has 11 clusters that include more than 2, 000 CORD-19 publications each (Figure A.5b). In total, these 11 clusters cover 54% of the CORD-19 publications. There are 90 clusters that include between 1, 000 and 2, 000, covering an additional 41% of the CORD-19 publications. The remaining 5% of the publications are scattered across 2, 429 small clusters each including fewer than 1, 000 publications. Of these clusters, 2, 012 consist of only a single publication.

To better understand the substantive focus of the different clusters, we overlay the clusters on the term map that we constructed previously. For each term in the term map, we selected all publications in which the term occurs and we calculated the percentage of these publications that belong to a specific cluster. This percentage determines the color of the term, with low percentages resulting in a blue color and high percentages resulting in a yellow color. For different clusters, different parts of the term map are highlighted in yellow, showing how clusters differ in their focus (Figure 7).

**Figure 7:**
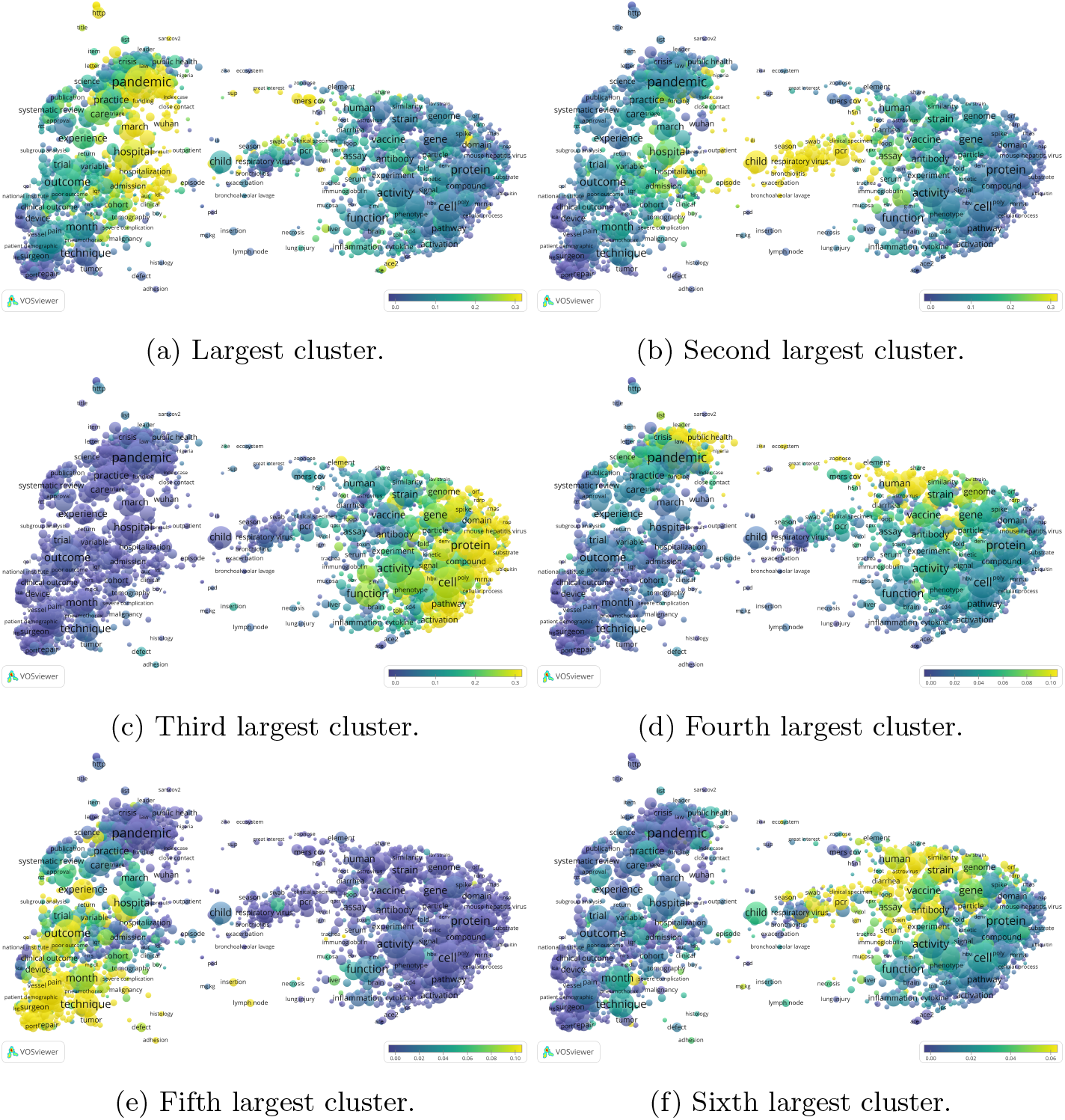
Bottom level citation network clustering. The color of a term reflects the percentage of publications in which the term occurs that belong to a specific cluster. For example, in (a), 39% of the publications that include the term “pandemic” belong to the largest cluster, whereas only 4% of the publications that include the term “protein” belong to this cluster. Compare with Figure A.7.

The largest cluster, consisting of 16, 913 publications, mainly covers publications dealing with the COVID-19 pandemic, including 15, 578 publications in 2020 and only about 100–200 in the preceding years. This focus is also clearly visible in the term map (Figure 7a). The second largest cluster, consisting of 10, 169 publications, focuses on respiratory viruses more generally (Figure 7b), showing a clear increase after the SARS epidemic in 2004, with again a clear increase in 2020. The third largest cluster, consisting of 7, 345 publications, has a clear biochemical focus, dealing with various proteins, transcriptions and pathways (Figure 7c) and shows a sustained increase from the early 2000s until 2020. The fourth largest cluster, consisting of 4, 375 publications, deals with both emerging infectious diseases and zoonosis (Figure 7d) and shows a similarly sustained increase from the early 2000s onward. The fifth largest cluster, consisting of 3, 252 publications, focuses on surgical procedures (Figure 7e) and shows an increase from 2007–2011 and then gently levels off towards 2020. Finally, the sixth largest cluster deals with viruses in animals (Figure 7f) and shows a longer history, dating back to the 1908s, with a gradual growth over the years and a steeper increase during the 2000s. The interpretation of these clusters is largely substantiated by comparing the clustering results to the topic model.

These relatively detailed clusters are hierarchically clustered at the top level. The largest cluster at the top level contains 20, 300 publications. It consists mainly of a combination of clusters focusing on the COVID-19 pandemic, among which the largest cluster at the bottom level (Figure 8a). The second largest cluster at the top level contains 13, 774 publications. It includes the third largest cluster and some smaller clusters at the bottom level, with a clear molecular biology focus (Figure 8b). The third largest cluster at the top level consists almost entirely of the second largest cluster at the bottom level, combined with a few smaller clusters. It has a focus on respiratory viruses. The three largest clusters at the top level cover 43% of all publications.

**Figure 8:**
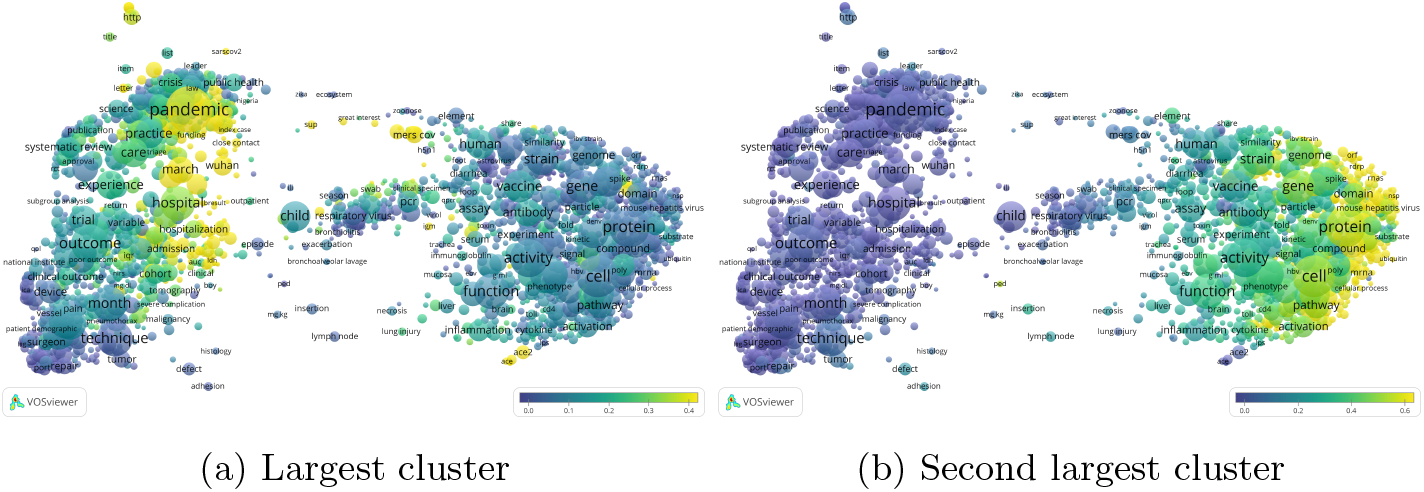
Top level citation network clustering. The color of a term reflects the percentage of publications in which the term occurs that belong to a specific cluster. For example, in (a), 46% of the publications that include the term “pandemic” belong to the largest cluster, whereas only 6% of the publications that include the term “protein” belong to this cluster. In (b), only 2% of the publications that include the term “pandemic” belong to the second largest cluster, whereas 54% of the publications that include the term “protein” belong to this cluster. Compare with Figure A.6.

We conclude this section by connecting our citation-based clustering results with our text-based topic modelling results. We use topic intensity to further qualify the results of the citation network clustering. More specifically, we characterise clusters by the average topic intensity of the publications they contain. We start with the top level clustering, and focus on the two largest clusters. The average topic intensities (Figure A.6) confirm that the first cluster contains publications focused on coronavirus outbreaks (COVID-19 in particular), including public health and clinical medicine research. The second cluster, on the other hand, has a clear molecular biology and immunology focus. Next, we consider the six largest clusters in the bottom level clustering (Figure A.7). While the third and sixth clusters are focused on molecular biology and immunology, the first, second and fourth are mainly concerned with coronavirus outbreaks, and the fifth covers clinical medicine.

Our citation-based results substantiate our earlier findings based on textual analyses. Both point to the existence of distinct research areas covered by CORD-19: coronaviruses, molecular biology research on viruses, public health and epidemics and other related topics (immunology, clinical medicine). These areas of research are interrelated, yet also contain specialised information, and they show different temporal trends.

### Altmetrics

CORD-19 publications have also been explored using Altmetric data, with the aim of describing their reception on social media, paying special attention to the dissemination of the publications across various social media sources. As can be seen in Table 1, a total of 88, 570 publications in the CORD-19 dataset (63%) have received some mention in Altmetric. This is a rather high coverage of publications compared to previous studies [17], which reported an overall coverage of 21.5% of 2012 publications on Twitter, and about 31% for publications in the biomedical and health sciences. This high coverage is even higher when the focus is on the most recent publications (i.e., those published in the early months of 2020), of which over 68% have received some social media mention covered by Altmetric.

**Table 1:** Coverage of CORD-19 publications by Altmetric.

|  | **CORD-19 publications** | **In Altmetric** | **Share in Altmetric** |
| --- | --- | --- | --- |
| **All** | 140,302 | 88,570 | 63.1% |
| **2020** | 52,440 | 36,058 | 68.8% |

Table 2 presents a more detailed description of the type of social media events around CORD-19 publications. We selected some of the most relevant sources covered by Altmetric, namely Twitter, blogs, recommendations in F1000Prime, news media mentions, citations in policy documents and citations in Wikipedia entries. Clearly, the most important source is Twitter, which by far accounts for the largest share of all (social) media interactions analysed. The second most important source are mentions in news media and blogs. The observation that there are more mentions in news media than in blogs also contrasts with previous studies [17]. This may signal the particular relevance of CORD-19 publications for mainstream news media. Another noteworthy characteristic is the recency of the publications being mentioned, particularly on Twitter and in news media. About 87% of all Twitter mentions relate to publications from 2020, and 75% of all mentions in mainstream news media also relate to 2020 publications.

**Table 2:**
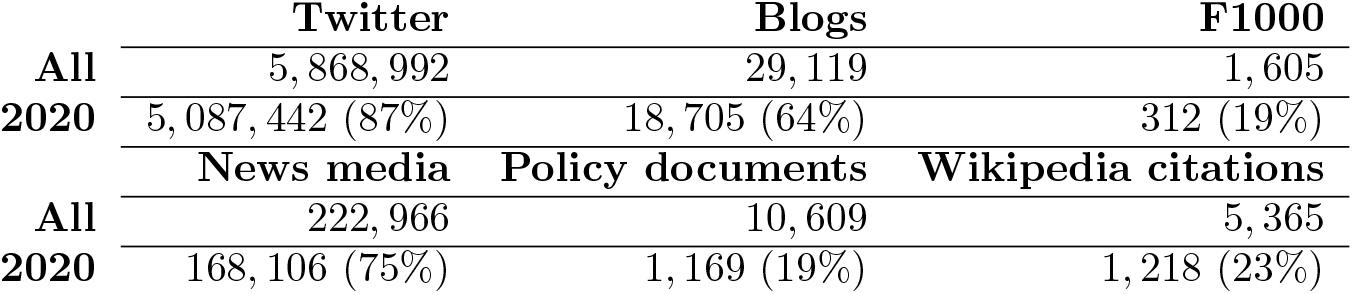
Social media events around CORD-19 publications.

|  | **Twitter** | **Blogs** | **F1000** |
| --- | --- | --- | --- |
| **All** | 5,868,992 | 29,119 | 1,605 |
| **2020** | 5,087,442 (87%) | 18,705 (64%) | 312 (19%) |
|  | **News media** | **Policy documents** | **Wikipedia citations** |
| **All** | 222,966 | 10,609 | 5,365 |
| **2020** | 168,106 (75%) | 1,169 (19%) | 1,218 (23%) |

Table 3 shows the main altmetrics per publication type. All CORD-19 publications with Altmetric data were matched with publications in the WoS database. In this way, a publication type was obtained for 82, 083 publications. The publication type classification of the WoS database was used because WoS offers a more detailed classification than databases such as Dimensions and Microsoft Academic [35].

**Table 3:** Main altmetrics per WoS publication type.

| Type | Publications | Twitter | News media | Blogs | Wikipedia |
| --- | --- | --- | --- | --- | --- |
| Article | 57,383 (70%) | 1,401,325 (46%) | 86,038 (63%) | 11,970 (63%) | 2,561 (56%) |
| Review | 11,217 (14%) | 288,135 (9%) | 14,256 (10%) | 2,451 (13%) | 1,394 (31%) |
| Editorial | 6,952 (8%) | 706,524 (23%) | 19,399 (14%) | 2,707 (14%) | 338 (7%) |
| Letter | 4,195 (5%) | 506,307 (17%) | 15,613 (11%) | 1,488 (8%) | 123 (3%) |
| News | 963 (1%) | 35,626 (4%) | 2,091 (2%) | 466 (2%) | 112 (3%) |
| Other | 1,373 (2%) | 4,136 (<1%) | 218 (<1%) | 54 (<1%) | 13 (<1%) |

While most publications in CORD-19 are classified as article (70%) or review article (14%), it turns out that editorials, letters and news items receive a disproportionate amount of attention on social and online media. As a reference, we also show the proportion of Wikipedia citations per publication type, where most attention turns out to be given to review articles. These findings are in line with previous work [17]. Since editorials, letters and news items usually present news, debates, and opinions, often using a less technical language than regular articles, it is not surprising that they are more appealing for social media. However, in CORD-19, these three publication types receive many more tweets (44% of all tweets) than what was previously found (below 20%) [17].

Figure 9 presents term maps showing the altmetrics reception of CORD19 publications. We cover the most immediate social media sources (or ‘fast’ sources [14]), which provide the earliest signals of the reception of publications. Twitter, blogs and news media all present a similar pattern, with a strong orientation towards the most recent COVID-19 publications, well captured by the largest cluster in both the bottom level and the top level citation network clustering (see Figures 7a and 8a). In terms of topics, terms such as ‘pandemic’, ‘hospital’, ‘hospitalization’ are among the most common terms used in publications mentioned in these social media platforms. These results are in line with those discussed in [15].

**Figure 9:**
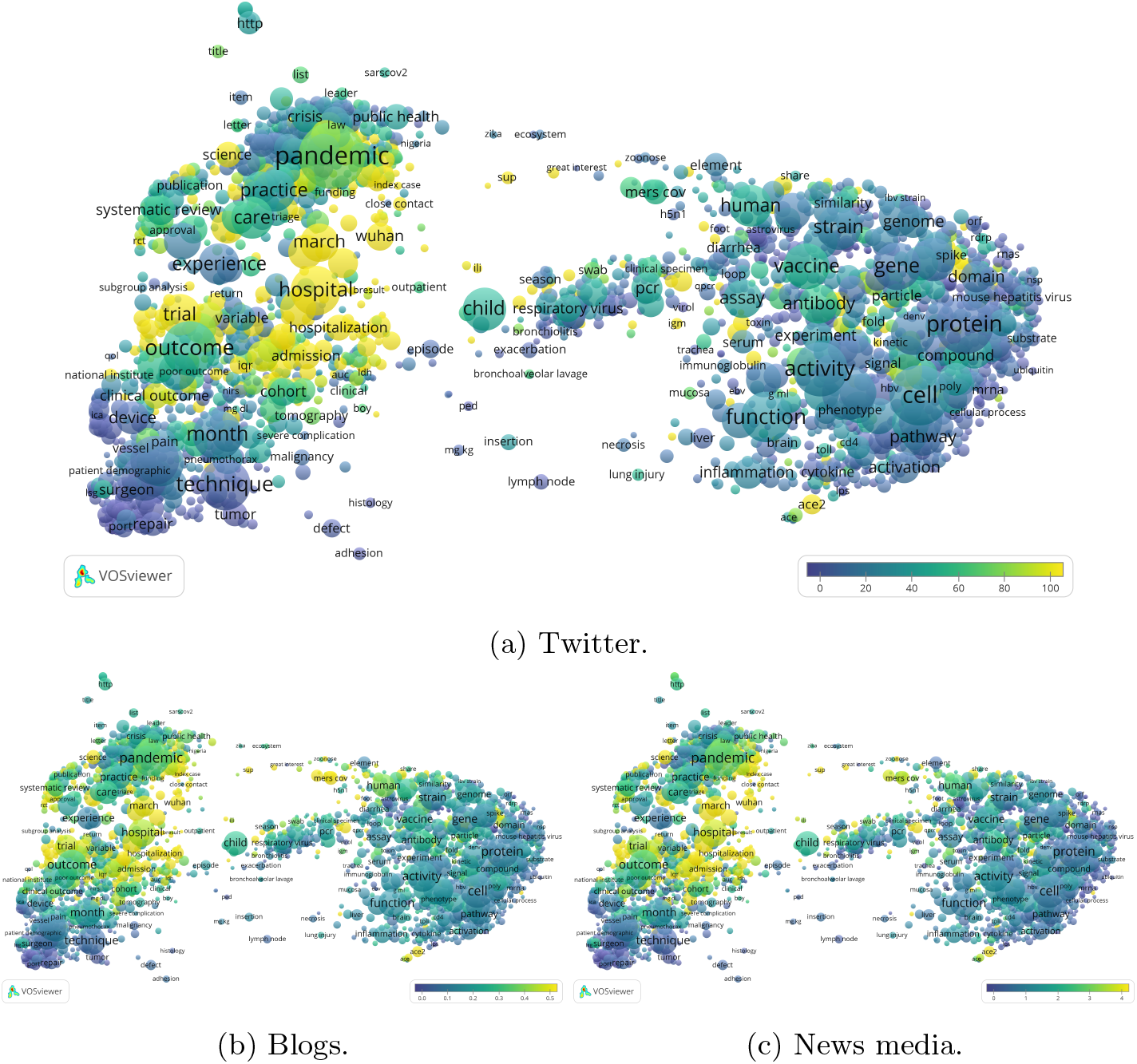
Immediate altmetric sources: Twitter, blogs and news media.

This preliminary altmetric analysis shows a strong present-day attention for research covered by the CORD-19 dataset. The majority of the tweets, blogs and news media mentions are focused on research produced during the current COVID-19 pandemic. The publication types that, in relative terms, receive most attention are editorials, letters and news items. This demonstrates the important role of social media, especially Twitter, in discussing COVID-19 research [22]. During a global pandemic like COVID-19, research is subject to high degrees of uncertainty [13, 36]. Rapidly increasing levels of social media activity around topics on which there is little academic consensus may increase the risk of scientific advise being misunderstood or misused by substantial segments of society. Social media analysis can provide tools for identifying and characterising areas with high levels of social media activity that may be in dissonance with the academic discourse. In future work, we plan to find and characterise these areas of discrepancy in order to inform scientists, science communicators, journalists and the public at large.

## Conclusion

We performed an in-depth analysis of the CORD-19 dataset of publications on COVID-19 and coronavirus research [38]. Our analysis of the delineation of CORD-19 indicates that the dataset is broader than just COVID-19 and coronavirus research. On the other hand, when compared to the Web of Science database, it turns out that CORD-19 largely covers all literature directly focused on COVID-19 and coronaviruses.

We carried out a deeper analyses of CORD-19 in various ways. We created a map of relevant terms extracted from the titles and abstracts of COVID-19 publications. This map confirmed the broad content of the dataset. A topic modelling analysis showed that CORD-19 publications are related to biomedical research on viruses in a broad sense, which includes research on COVID-19 and coronaviruses. Dominant topics in CORD-19 also include research on public health and epidemics; molecular biology; other coronaviruses; influenza and other families of viruses; immunology and antivirals; methodology (testing, diagnosing, trials). Furthermore, the topic intensity over time is far from uniform. Coronavirus research has followed known outbreaks (SARS, MERS, COVID-19) and for years before 2020 this research represents only a small portion of CORD-19.

We performed a citation network clustering analysis using data from Dimensions. Citation network clusters highlighted the relative cohesiveness of CORD-19. In line with the textual analyses, the clusters confirmed the broad coverage of the dataset. Overall, there seem to be two prominent citation clusters: one, more recent, that covers research on specific coronaviruses, with a public health and epidemiological focus, and another one, with a longer time span, focused on molecular biology. Molecular biology research on viruses is, in general, a prominent component of CORD-19. It is likely that, as research on COVID-19 rapidly expands, these themes will broaden as well.

Lastly, we considered Altmetric data, in order to gauge how much attention CORD-19 research has attracted over time. The current COVID-19 outbreak dominates attention on social media, in particular from Twitter, highlighting the interest for scientific results during the COVID-19 pandemic. Editorials and letters in CORD-19 get a disproportionate amount of attention on social media. Our work acknowledges that research on viruses, and coronaviruses specifically, does not exist in a vacuum. Delimiting research on a certain subject matter requires difficult choices that inevitably involve a certain degree of arbitrariness. We praise the breadth and relative coherence of CORD-19. This dataset rightly merits attention and is useful to allow many researchers to engage with the topic. Nevertheless, we also suggest that critical awareness is required when using CORD-19, as our results demonstrate that its contents cover a broad set of topics. Different subsets of CORD-19 should be used for specific purposes, for example for making historical analyses on funding of specific research topics or for automatically extracting structured information. We exemplified some approaches to segment CORD-19 in various ways.

Clearly, there are many areas for future COVID-19 work by the scientometric community. We conclude by suggesting three areas in particular. Firstly, there seems to be a need for a comprehensive and ongoing mapping of COVID-19-related research. A multidisciplinary map of COVID-19-related research, considering diverse disciplinary perspectives and information needs, will be useful to surface relevant research, also outside the biomedical domain. Secondly, CORD-19 provides a virtuous example of open data sharing. The scientometric community can contribute by creating and maintaining additional datasets on COVID-19 research. Thirdly, as shown by our results, there is a lot of social media attention for COVID-19 research. Indeed, the role of information, and especially reliable scientific information, has been central to the unfolding of the current pandemic [41]. Consequently, a relevant area for future work is to better understand the mechanics of online scientific information diffusion, using altmetrics and other data sources. This line of work has the potential to provide valuable information to experts and governments during the current and future pandemics.

## Data availability

Most of our analysis can be replicated using the code and following the instructions given in the accompanying repository: https://github.com/CWTSLeiden/cwts_covid. We welcome contributions and suggestions, ideally by opening an issue or doing a pull request. Analyses based on Altmetric, Dimensions, Twitter and Web of Science data require access to these services.

## Acknowledgements

We thank Digital Science for providing access to Altmetric and Dimensions data and for constant support throughout our project. We also thank our colleagues at CWTS for enthusiastic and insightful discussions on how the science studies community can contribute to research on COVID-19. We particularly thank Zhichao Fang and Jonathan Dudek for their help in the data collection for this study.

## A Appendix

## A.1 Topic top words

Top 20 words per topic, using the LDA model. Words consisting of one or two characters are filtered out. Compare with Figure A.4 for the topic intensity over time.

- **Topic 0, Public health**: “respiratory”, “infection”, “virus”, “patient”, “child”, “influenza”, “acute”, “clinical”, “viral”, “pneumonia”, “symptom”, “test”, “diagnosis”, “case”, “positive”, “detect”, “tract”, “severe”, “cause”, “study”.
- **Topic 1, Coronavirus outbreaks**: “covid-19”, “COVID-19”, “pandemic”, “health”, “sars-cov-2”, “coronavirus”, “2020”, “public”, “care”, “2019”, “patient”, “covid-19 pandemic”, “hospital”, “medical”, “emergency”, “lockdown”, “healthcare”, “public health”, “response”, “Health”.
- **Topic 2, Clinical medicine**: “group”, “study”, “compare”, “rate”, “significantly”, “high”, “result”, “patient”, “conclusion”, “year”, “difference”, “analysis”, “control”, “score”, “significant”, “respectively”, “method”, “associate”, “mean”, “total”.
- **Topic 3, Molecular biology**: “virus”, “sequence”, “coronavirus”, “strain”, “gene”, “assay”, “sample”, “antibody”, “isolate”, “detect”, “analysis”, “study”, “protein”, “human”, “genome”, “calf”, “result”, “acid”, “detection”, “infectious”.
- **Topic 4, Public health**: “study”, “review”, “include”, “health”, “result”, “datum”, “report”, “intervention”, “search”, “evidence”, “method”, “practice”, “quality”, “research”, “risk”, “systematic”, “literature”, “conduct”, “identify”, “article”.
- **Topic 5, Coronavirus outbreaks**: “disease”, “case”, “infection”, “outbreak”, “SARS”, “transmission”, “country”, “epidemic”, “control”, “syndrome”, “respiratory syndrome”, “severe”, “report”, “severe acute”, “risk”, “respiratory”, “infectious”, “death”, “China”, “spread”.
- **Topic 6, Clinical medicine**: “patient”, “treatment”, “case”, “aneurysm”, “clinical”, “treat”, “lesion”, “artery”, “chest”, “stroke”, “image”, “acute”, “outcome”, “result”, “follow-up”, “imaging”, “occlusion”, “report”, “endovascular”, “complication”.
- **Topic 7, Clinical medicine**: “patient”, “surgery”, “laparoscopic”, “surgical”, “complication”, “procedure”, “undergo”, “postoperative”, “perform”, “technique”, “case”, “result”, “time”, “pain”, “group”, “method”, “repair”, “resection”, “patient undergo”, “mean”.
- **Topic 8, Immunology**: “cell”, “infection”, “response”, “expression”, “mouse”, “disease”, “lung”, “immune”, “increase”, “role”, “effect”, “gene”, “receptor”, “study”, “tissue”, “mechanism”, “induce”, “level”, “cytokine”, “function”.
- **Topic 9, Epidemics**: “model”, “datum”, “number”, “analysis”, “time”, “result”, “network”, “different”, “base”, “method”, “dynamic”, “study”, “propose”, “social”, “epidemic”, “paper”, “approach”, “individual”, “population”, “estimate”.
- **Topic 10, Epidemics**: “method”, “result”, “study”, “device”, “pressure”, “test”, “image”, “temperature”, “flow”, “evaluate”, “forecast”, “compare”, “measure”, “high”, “tissue”, “increase”, “system”, “concentration”, “performance”, “time”.
- **Topic 11, Molecular biology**: “protein”, “virus”, “viral”, “cell”, “replication”, “coronavirus”, “activity”, “antiviral”, “membrane”, “sars-cov”, “domain”, “infection”, “structure”, “host”, “binding”, “inhibitor”, “fusion”, “interaction”, “hepatitis”, “site”.
- **Topic 12, Immunology**: “cell”, “vaccine”, “antibody”, “mouse”, “response”, “infection”, “immune”, “antigen”, “induce”, “virus”, “culture”, “human”, “line”, “t cell”, “vitro”, “vaccination”, “immunity”, “in vitro”, “challenge”, “recombinant”.
- **Topic 13, Public health**: “disease”, “drug”, “development”, “review”, “human”, “research”, “treatment”, “potential”, “approach”, “system”, “provide”, “clinical”, “recent”, “include”, “develop”, “discuss”, “current”, “technology”, “application”, “pathogen”.
- **Topic 14, Clinical medicine**: “patient”, “treatment”, “clinical”, “severe”, “blood”, “disease”, “therapy”, “level”, “cancer”, “care”, “intensive”, “serum”, “high”, “failure”, “plasma”, “oxygen”, “risk”, “unit”, “heart”, “outcome”.

**Figure A.1:**
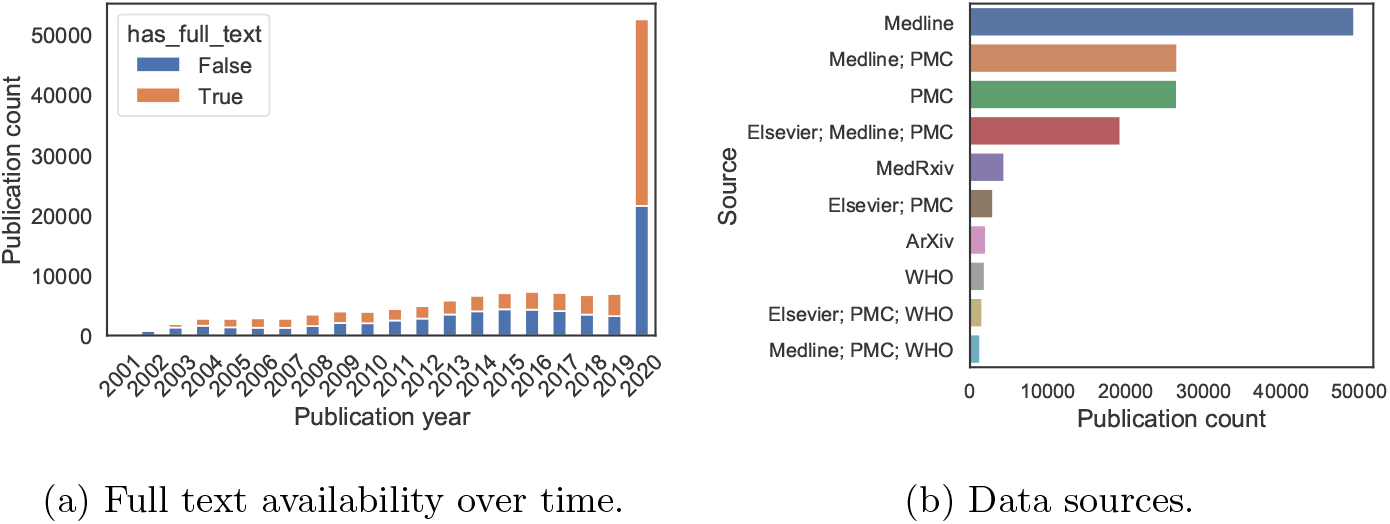
Full text availability and data sources.

**Figure A.2:**
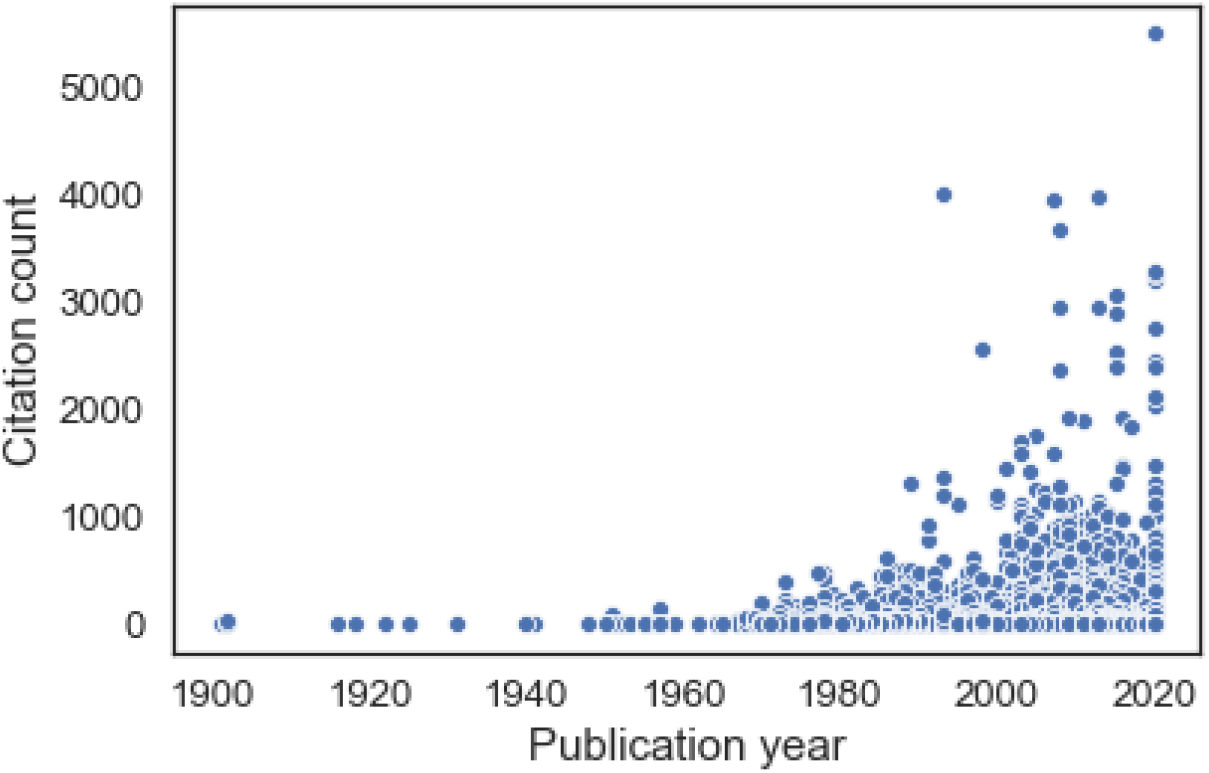
Scatter plot of the number of citations received by articles from different years.

**Figure A.3:**
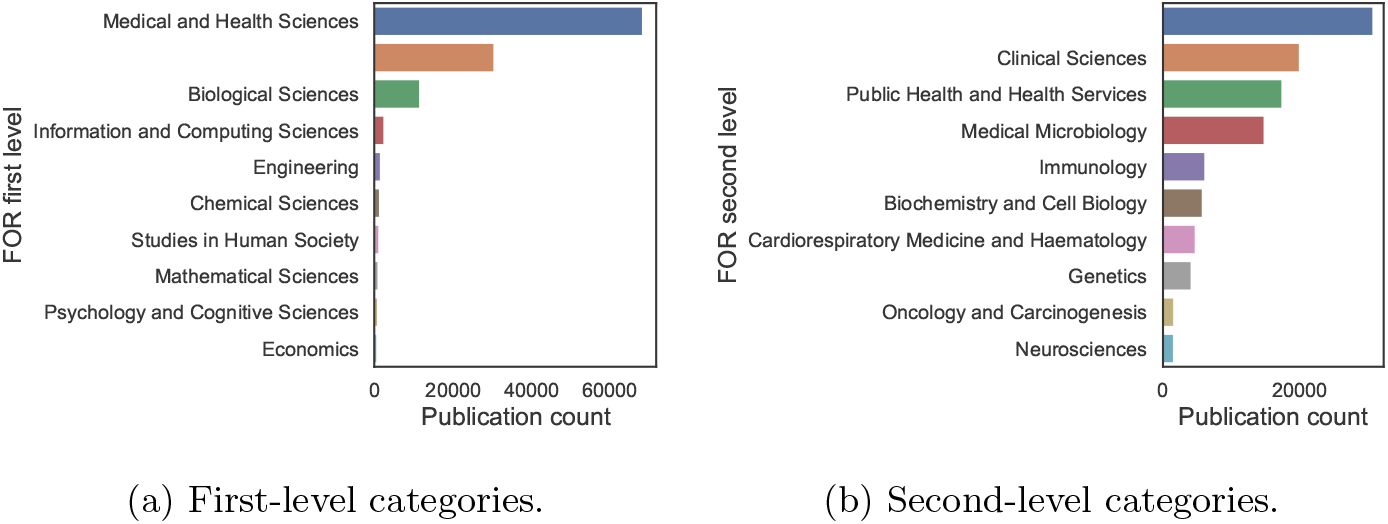
Categories in the FOR classification in Dimensions. The empty label accounts for articles without a FOR category.

**Figure A.4:**
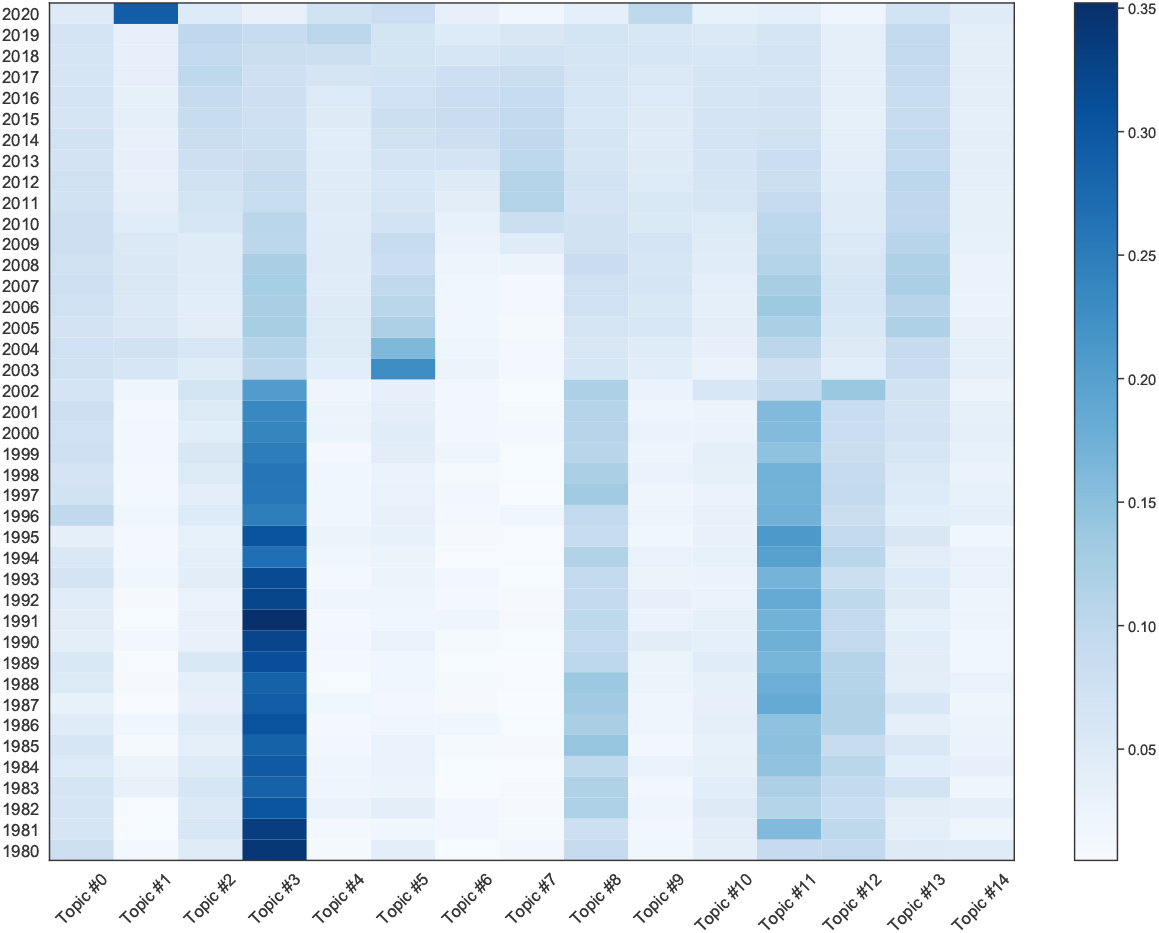
Topic intensity over time, using the LDA model. (a) Top level clustering: fewer, larger clus-(b) Bottom level clustering: more, smaller ters. clusters.

**Figure A.5:**
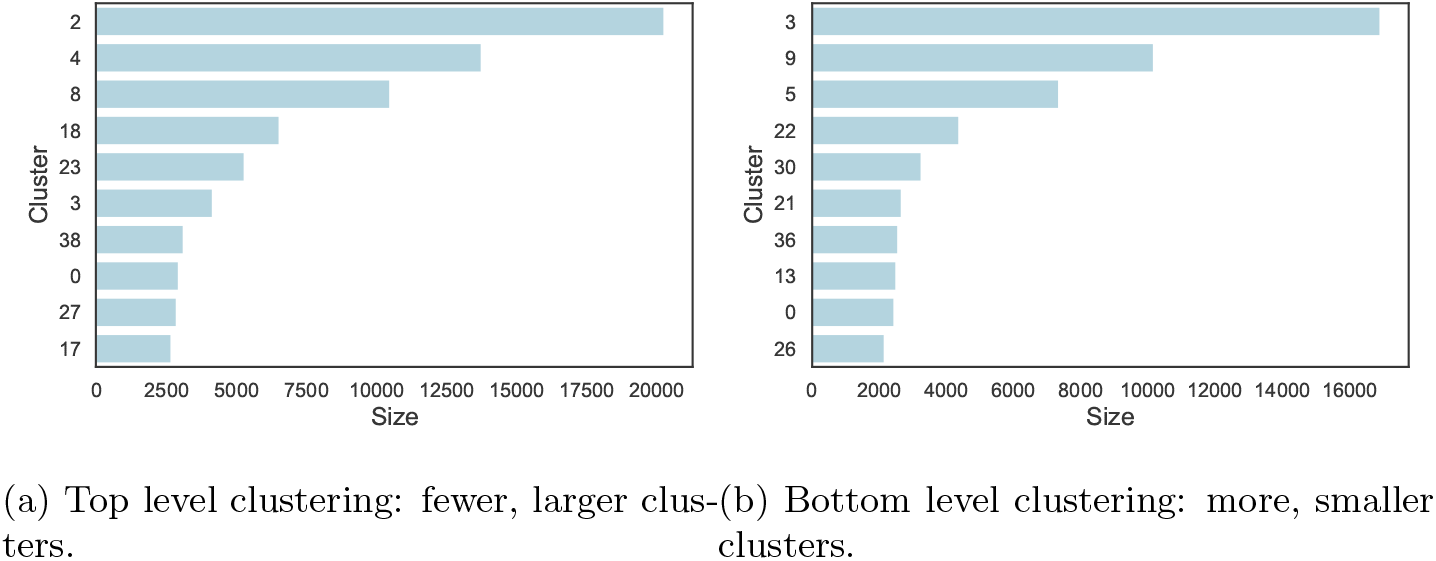
Cluster size in the top level and the bottom level clustering of the citation network.

**Figure A.6:**
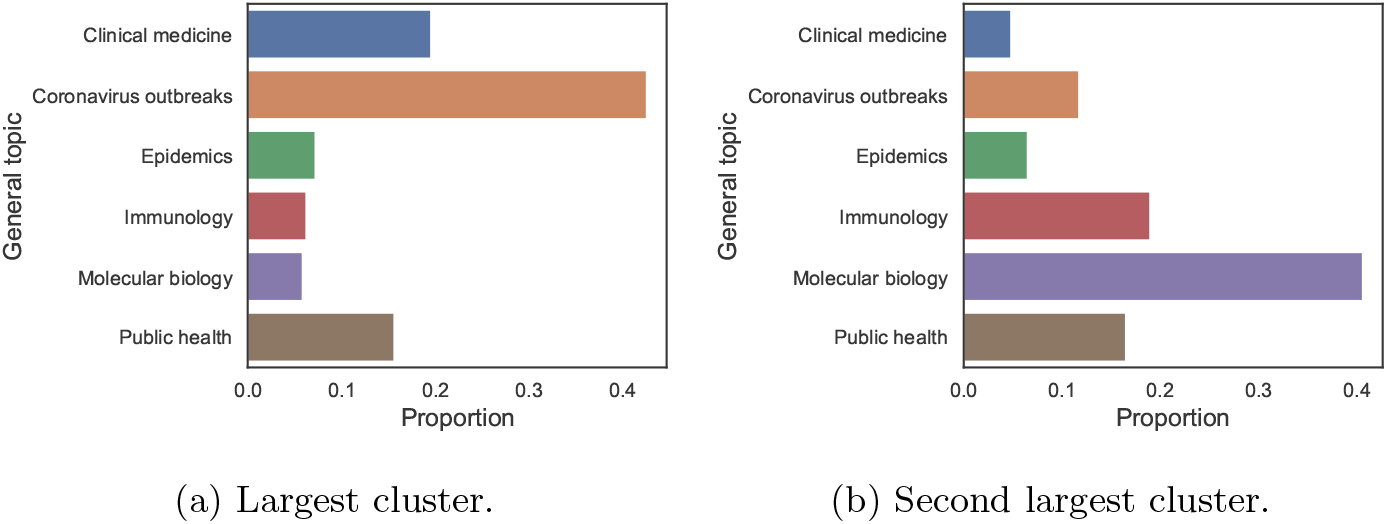
Breakdown by topic for the top level clustering of the citation network.

**Figure A.7:**
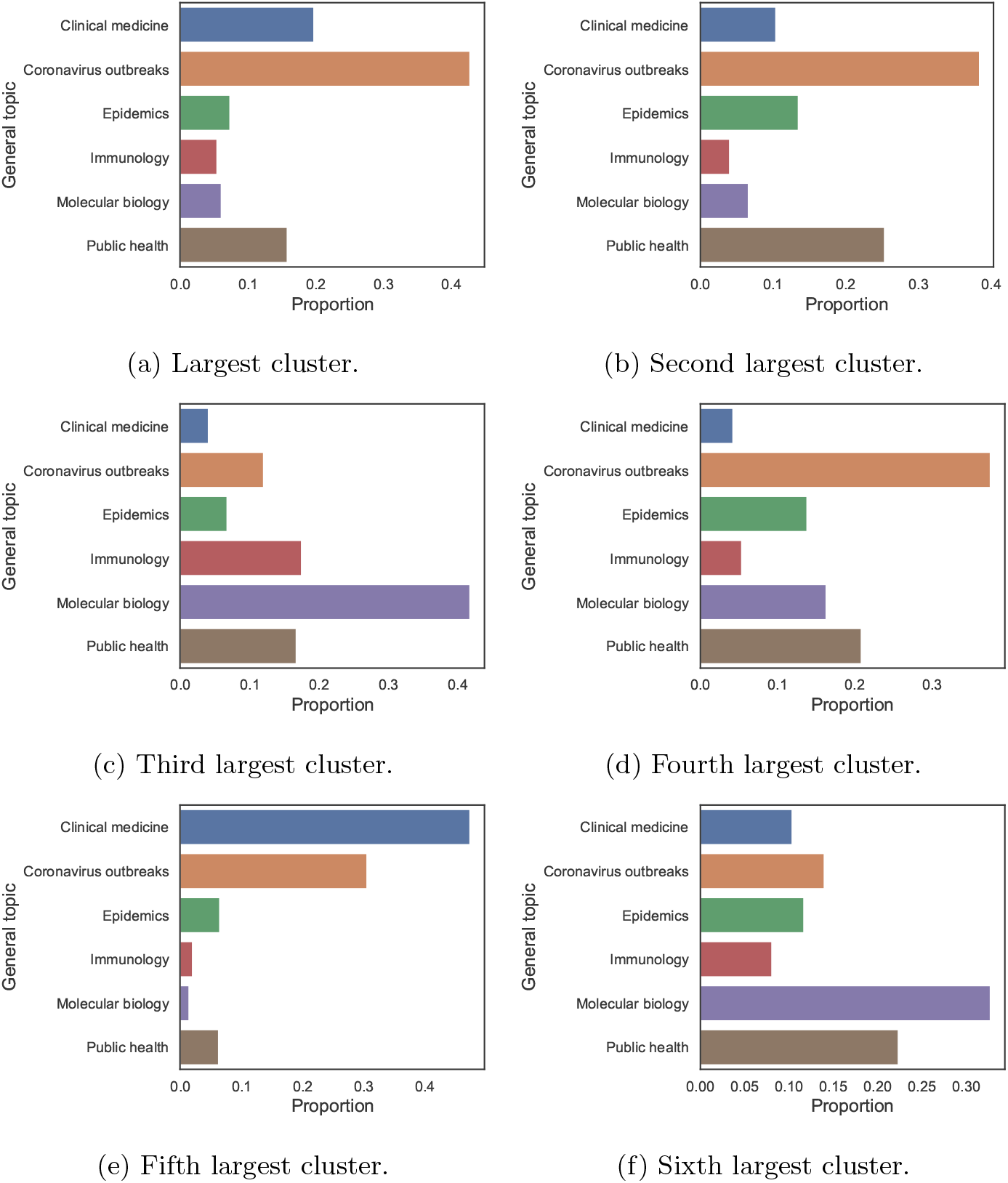
Breakdown by topic for the bottom level clustering of the citation network.

## Footnotes

1 For coronaviruses, see [16].

2 This call has been taken up: For example, the ACL conference has run an emergency NLP COVID-19 workshop which mentioned the CORD-19 dataset on its call for papers (https://www.nlpcovid19workshop.org) and a TREC-COVID challenge has been announced on CORD-19 (https://ir.nist.gov/covidSubmit).

3 https://app.dimensions.ai/browse/categories/publication/for.

4 Elsewhere in this paper, we use data from Dimensions instead of WoS. Because we do not have access to abstracts in Dimensions, we use WoS data in this section.

5 Using tomotopy, https://bab2min.github.io/tomotopy [version 0.7.0].

6 We used the Leiden algorithm implementation from igraph.

